# An Ultrasensitive Biosensor for Quantifying the Interaction of SARS-CoV-2 and Its Receptor ACE2 in Cells and *in vitro*

**DOI:** 10.1101/2020.12.29.424698

**Authors:** Xiaolong Yang, Lidong Liu, Yawei Hao, Yee Wah So, Sahar Sarmasti Emami, Derek Zhang, Yanping Gong, Prameet M. Sheth, Yu Tian Wang

## Abstract

The severe acute respiratory syndrome coronavirus-2 (SARS-CoV-2) is currently spreading and mutating with increasing speed worldwide. Therefore, there is an urgent need for a simple, sensitive, and high-throughput (HTP) assay to quantify virus-host interaction in order to quickly evaluate infectious ability of mutant virus and develop or validate virus-inhibiting drugs. Here we have developed an ultrasensitive bioluminescent biosensor to evaluate virus-cell interaction by quantifying the interaction between SARS-CoV-2 receptor binding domain (RBD) and its cellular receptor angiotensin-converting enzyme 2 (ACE2) both in living cells and *in vitro*. We have successfully used this novel biosensor to analyze SARS-CoV-2 RBD mutants, and evaluated candidate small molecules (SMs), antibodies, and peptides that may block RBD:ACE2 interaction. This simple, rapid and HTP biosensor tool will significantly expedite detection of viral mutants and anti-COVID-19 drug discovery processes.

---

The recent outbreak of coronavirus SARS-CoV-2 causes human respiratory disease called COVID-19 in China and worldwide. At the time of writing this manuscript, over 40 million cases and 1 million deaths have been attributed to the virus. The key in the fight against this virus is the development of fast, accurate, sensitive diagnostic tools, and potent therapeutic drugs. Most recently, multiple tools, which include fast detection of viral mRNAs and anti-viral antibodies (IgM and IgG) in COVID19+ patients, have been developed^1^. However, drugs effectively eradicating the virus are still lacking. Currently, the most promising approach to stop the virus is to develop vaccine against this virus. However, vaccine development and testing usually require long time (>12 months). In addition, as for vaccines against other types of virus such as SARS, there is no guarantee that the vaccine will work. Moreover, given the nature of this SARS-CoV-2 virus, it is also possible that it will become endemic and circulate in the population like seasonal influenza virus (flu). And, like the flu, it could mutate into new strains that escape the vaccines. Therefore, for the long-term fight against COVID-19, other long-lasting therapeutic drugs targeting this virus are also required.

Beside vaccines, two major approaches have been proposed or reported to target the SARS-CoV-2^2^: 1) Use of SMs, antibodies or peptides to inactivate proteins involved in viral entrance [e.g. viral Spike (S) protein (anti-S antibodies)], processing [e.g. cleavage by proteases M^pro^ and 3CL^pro^ (Lopinavir-ritonavir), and endocytosis (e.g. hydroxychloroquine)], and replication [e.g. RNA polymerase (remdesivir)]; 2) Use of convalescent plasma/sera containing anti-SARS-CoV-2 antibodies from recovered patients. However, up until now, there are no developed drugs used in clinical treatment that can effectively treat COVID-19 and resulting in reduced mortality. Since clinical trials usually take years to accomplish, there is an urgent need for tools that can quickly test the effectiveness of candidate anti-virus drugs to expedite the discovery processes. In addition, although many SARS-CoV-2 mutants have been detected in new COVID-19 patients globally^3^, there is no simple, fast and accurate way to test whether these mutants will affect their entrance into cells via changed interaction with ACE2 on the cell membrane.

It has been reported that SARS-CoV-2 enters human epithelial cells by using the RBD of its S protein to attach to a receptor called ACE2 protein on the cell surface^4^. Therefore, blocking RBD:ACE2 protein-protein interaction (PPI) will be an excellent strategy to curb virus entrance/invasion into cells and their subsequent infection^5^. The gold standard to test the effect of RBD/ACE2 mutation, drugs or antibodies on their interactions is to use viral neutralization assays using either live virus^6^ or viral vectors pseudotyped with the S protein^7^. However, these *in vivo* virus-based assays require a level III biosafety facility and are time-consuming to run. Although some *in vitro* protein-based assays such as surface plasmon resonance (SPR)^8^ and ELISA^9–11^ have also been developed in analyzing RBD:ACE2 interactions, they are less sensitive (color detection) and technically challenging (require purification of proteins from mammalian cells and multiple steps or special equipment to perform), which makes them difficult to operationalize in most research and clinical labs. Therefore, a simple, sensitive, and fast assay that can be used to perform HTP screening or validation for drugs blocking RBD-ACE2 PPI for anti-COVID-19 therapy is required.

In 2016, an ultrasensitive split luciferase complementary assay (SLCA) called NanoLuc Binary Technology (NanoBiT) was developed to monitor PPIs^12^. This assay is based on the observation that the NanoLuc luciferase enzyme can be separated into two fragments, 18 kDa Large BiT (LgBiT or Lg) and 11 amino acids (aa) small BiT (SmBiT or Sm) with each fragment fused to a target protein of interest. PPI of these two proteins can lead to reconstitution of active NanoLuc luciferase from the LgBiT and SmBiT and emit light in the presence of its substrate furimazine. By using this technology, we successfully constructed a NanoBiT YAP:TEAD biosensor and successfully used it to screen and identify SMs disrupting YAP-TEAD PPIs in cancer therapy^13^. In this study, we have used this simple and powerful technology and developed the first NanoBiT biosensor to quantify SARS-CoV-2 RBD:ACE2 PPI both *in vitro* and *in vivo*. We have also validated this biosensor to test the effects of SMs, antibodies, and peptides on RBD:ACE2 PPIs.

## MATERIALS AND METHODS

### Materials

Chemicals were purchased from Sigma-Aldrich or Bioshop unless otherwise stated.

### Biosensor plasmid construction and *in vitro* mutagenesis

To construct <u>S</u>ARS-CoV-2 <u>R</u>BD:<u>A</u>C<u>E2</u> biosensor (SRAE2-BS), RBD (aa. 319-685) or ACE2 ectodomain (aa. 18-615) was first amplified from by PCR from codon optimized SARS-Cov-2 (2018-nCoV) Spike S1 ORF (SinoBiologicals Inc.) or human ACE2 cDNAs (SinoBiologicals Inc.; accession number NM_021804.1), respectively, using PrimStar DNA polymeras (Takara Bio). LgBiT, SmBiT, GS linker (GSSGGGGSGGGGSSG), or 6×His tag (Fig. 1A) was added into either N- or C-terminal PCR products of the RBD or ACE2 by including their sequences in primers or by overlapping PCR (see Table 1S for primer sequences). The amplified PCR products were digested with EcoRV/NicoI and subcloned into pFUSE_hIgG1_Fc2 mammalian expression vector (Invivogen), which contains a N-terminal IL2 secretion signal peptides (SP).

**Figure 1.**
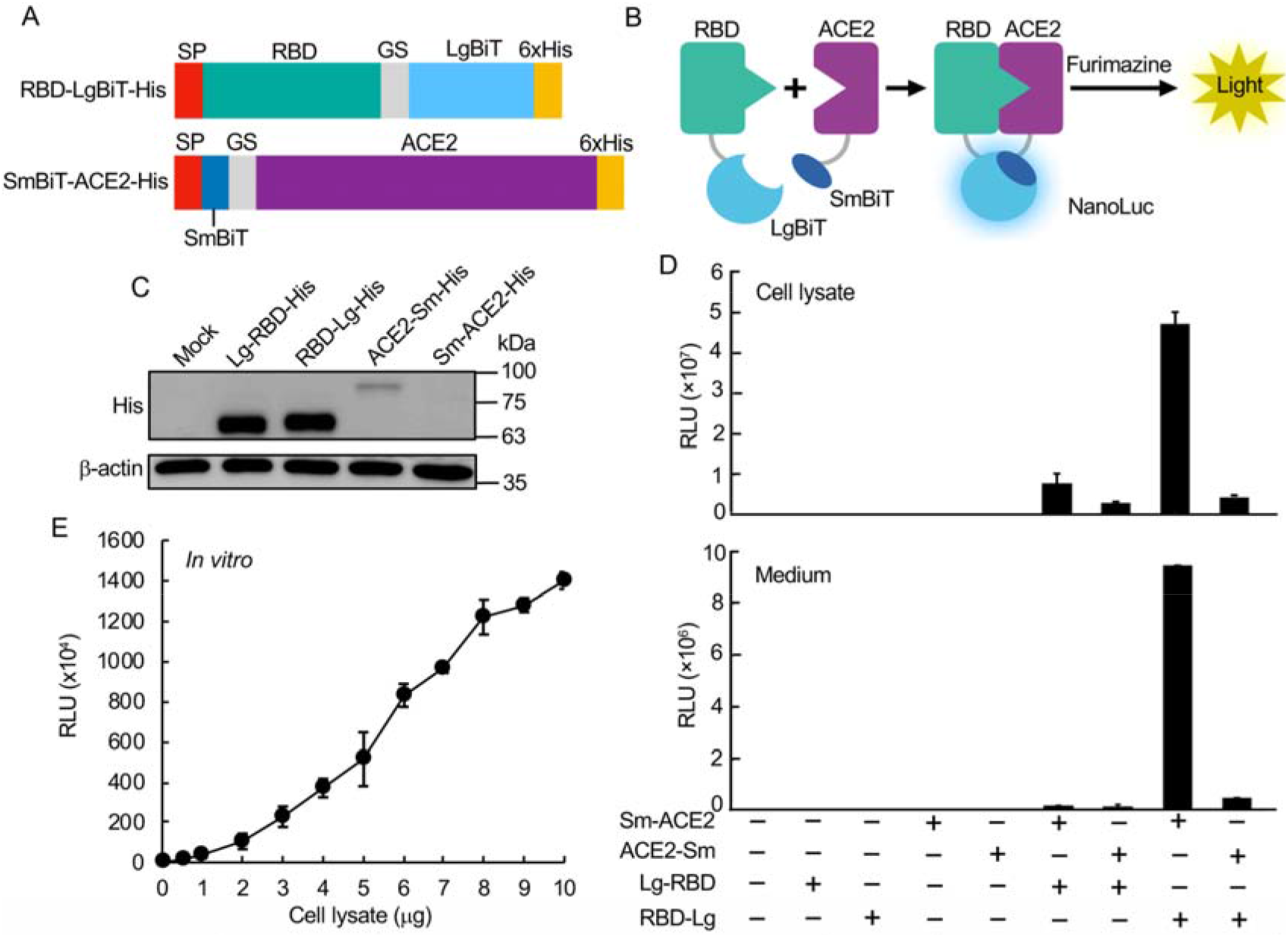
Construction and validation of RBD:ACE2 NanoBiT biosensor. A. Schematic of SRAE2-BS constructs. SP, signal peptide. B. Demonstration of SRAE2-BS working mechanism. C. Western blot analysis of protein expression of SRAE2 components. Plasmids were transfected into HEK293T cells in 12-well plate. 10 μg of protein lysates were subjected to SDS-PAGE, followed by western blot analysis using anti-His first antibody, ß-actin was used as internal protein loading control. Molecular weight in kilodalton (kDa) is shown on the right of the blot. D. Luciferase analysis of different combinations of biosensor components. Plasmids were transfected into HEK293T cells in 96-well plate, followed by luciferase analysis using cell lysates from proteins extracted from transfected cells (upper panel) or secreted protein from cell culture medium (lower panel). RLU, relative luciferase unit. E. Luciferase analysis of SRAE2-BS activity *in vitro*. Equal amounts (0-10 μg) of Sm-ACE2-His and RBD-Lg-His protein lysates were mixed and incubated together for 30 min, followed by luciferase analysis.

PCR was used to construct SRAE2-BS plasmids with ACE2 or RBD deletions, whereas overlapping PCR was used to make *in vitro* mutagenesis. The primer sequences were described in Table S2.

### Cell culture

HEK293T (human embryonic kidney) cells were cultured in Dulbecco’s modified Eagle’s medium (DMEM; Sigma D6429) containing 10% FBE, and 1% Penicillin/Streptomycin (Invitrogen) at 37 □ °C with 5% CO_2_.

### DNA Transfection and western blot analysis

About 1 μg/well of SRAE2-BS plasmid was transfected into HEK293T cells in 6-well plate using PolyJet transfection reagent (SignaGen). Two days after transfection, protein was extracted using RIPA lysis buffer (50 mM Tris.HCl, 150 mM NaCl, 0.02% sodium azide, 1% NP-40, 0.1% SDS, and 0.5% sodium deoxycholate). 10-20 μg extracted protein lysates were separated by 10% SDS-polyacrylamide gel electrophoresis (PAGE)(Bio-Rad) and transferred to nitrocellulose membranes. The blot was first probed with anti-6×His mouse monoclonal antibody (Abcam ab18184), followed by incubation with goat anti-mouse IgG secondary antibody (Jackson ImmunoResearch). The signal was detected using ECL chemiluminescence substrate and visualized using an Amersham Imager 600UV.

### Peptide design and synthesis

The interrupting peptides were designed based on the analysis of the binding sequences between the SARS-CoV2 spike glycoprotein and ACE2, particularly, the amino acid residues between 446 and 505 on RBD and residues 19-393 on ACE2, which was believed to form hydrogen bonds with ACE2 residues 19-393 at SARA-CoV-2 RBD/ACE-2 interface^8,14,15^. The final sequences of the modified peptides were optimized by using peptide-protein docking software, HPEPDOCK^16^ and HADDOCK^17^. The peptides with top docking energy scores were selected for the biosensor binding assay. Peptides were synthesized on a Liberty Blue microwave aided peptide synthesizer (CEM Corporation) using solid phase methodology and Fmoc-protecting group strategy on a Rink Amide MBHA resin (Gyrosprotein Technologies). In each peptide-build-up cycle, 20% piperidine in DMF (N,N-dimethylmethanamide) was used for N-terminal Fmoc-deprotection and N,N’-diisopropylcarbodiimide (DIC) /Oxyma were used as coupling reagents. Synthesized peptides were cleaved from resin in TFA cleavage cocktail containing trifluoroacetic acid, phenol, deionized water, thioanisole and 1,2-ethanedithiol (82.5:5:5:5:2.5 %, v/v) for 2 hours at room temperature and precipitated with ice-cooled methyl tert-butyl ether. Crude peptides were further purified on an Agilent 1200Preparative HPLC coupled with 6100 Mass Spectrometry system (Agilent) using a reverse phase C-18 column (ACE C18-300, 250×21.2mm, 10μm) with water/acetonitrile gradient elution. The final purity of all peptides is >95%.

### NanoLuc luciferase (NanoBiT) assay

For analysis of biosensor activity in living cell *in vivo*, 2×10^4^ HEK293T cells were seeded in triplicate in 96-well plate 24 hours before transfection. 100 ng of Lg-RBD, RBD-Lg, Sm-ACE2, or ACE2-Sm plasmids were transfected alone or together into HEK293T cells using PolyJet transfection reagent (SignaGen). 24 hours after transfection, the medium was transferred to new wells. The attached cells were lysed in 20 μl of 1× passive lysis buffer (1× PLB, Promega) at room temperature (RT) for 15 min. 20 μl of cell lysate or medium were subjected to NanoLuc luciferase assay using Nano-Glo Live Cell Reagent containing 1/50 diluted furimazine substrate (Promega). Relative Luminescence Unit (RLU) was measured using GloMax Navigator Microplate Luminomer (Promega).

For analysis of biosensor activity *in vitro*, 500 ng/well of wild-type (WT) or mutant (deletions or point mutations) Sm-ACE2 or RBD-Lg were transfected into 12-well plate using PolyJet transfection reagent (SignaGen). Two days after transfection, cells were lysed in 1× PLB at RT for 15 min. Protein concentration were quantified using RC DC Protein Assay kit (Bio-Rad). Protein lysates were diluted into 1 μg/μl and kept at −80 °C. For *in vitro* biosensor analysis, equal amounts of WT or mutant RBD-Lg and Sm-ACE2 were mixed together in 10 μl and incubated at RT for 30 min. 10 μl of 1/50 diluted furimazine substrate were added, followed by measurement of RLU using GloMax Microplate luminometer.

All experiments were repeated at least 2-3 times. The means and standard deviations (S.D.) of RLU of triplicate samples were shown.

### Validation of SRAE2-BS using SMs, antibodies, and peptides

About 6 μg of Sm-ACE2 or RBD-Lg plasmids were transfected into a 100 mm plate using PolyJet. Two days after transfection, protein lysates were extracted, quantified, diluted, and stored as described above. For examination of the effect of candidate drugs on biosensor activity, triplicate of increasing amounts (0-1000 μM) of SMs [Baicalin, Hesperidin, Sculellarin, Theaflavin, Heparin from TargetMol] or freshly prepared synthesized peptides [ACE2-P1~2] or 1-50 μg/ml of IgG or VHH72 anti-SARS-CoV-2 Spike RBD LIamabody monoclonal antibody (R&D#LMAB10541) were preincubated with 1 μg of Sm-ACE2 protein lysate in 10 μl in 96-well plates on a rocker at RT for 30 min. 1 μg/10 μl of Sm-ACE2 protein lysates were added and the mixture was incubated on a rocker at RT for another 30 min. For peptides targeting ACE2 (RBD-P1~3), increasing concentrations (1-100 μM) of peptides were preincubated with Sm-ACE2 protein lysate for 30 min, followed by incubation with RBD-Lg protein lysates. 10 μl of 1/50 diluted furimazine substrate were added into each well. The mixtures were incubated on a rocker for 2 min, followed by measurement of RLU using GloMax Microplate Luminometer (Promega). All experiments were repeated at least twice. The mean and standard error (S.E.) of triplicate samples were calculated.

### Statistical analysis

Student’s t-test (2-tailed) was used for statistical analysis. A value of P ≤ 0.05 was accepted as statistically significant.

## RESULTS AND DISCUSSION

To build the NanoBiT bioluminescent biosensor monitoring the RBD:ACE2 PPI, we first construct biosensor plasmids by fusing LgBiT or SmBiT with RBD or ACE2, respectively, at their N- or C-termini. Interaction of RBD and ACE2 will make complementation of LgBiT and SmBiT to form NanoLuc luciferase, which will emit light in the present of its substrate furimazine (Fig. 1B). Western blot analysis of each construct shows that while similarly high levels of Lg-RBD-His and RBD-Lg-His were expressed in cells, relatively low levels of Sm-ACE2-His and ACE2-Sm were expressed (Fig. 1C). Luciferase analysis of biosensor activity indicates that transfection of each component of the biosensor alone had little luciferase activity, whereas the combination of Sm-ACE2 and RBD-Lg obtained the highest activity in both protein lysate extracted from cells transfected with plasmids and collected cell culture medium (secreted biosensor)(Fig. 1D). To analyze the biosensor activity *in vitro*, we combined increasing amounts (0-10 μg) of Sm-ACE2 protein lysates with equal amounts of RBD-Lg lysates *in vitro* and incubated at RT for 30 min, followed by luciferase analysis. Significantly, combination of only 1 μg of protein lysate of Sm-ACE2 and RBD-Lg can obtain over 1× 10^5^ RLU. The signal intensity linearly increases with increasing amounts of cell lysates added (Fig. 1E). These data strongly suggest that we are able to quantify RBD-ACE2 PPI both *in vivo* in cells and *in vitro* using protein lysates extracted from plasmid-transfected cells. This newly developed bioluminescent biosensor tool is simple (only a luciferase assay *in vitro*), sensitive (~0.2-1 μg crude cell lysate per assay; no protein purification is required), fast (30 min *in vitro*), accurate (quantified by light emission unit), HTP (96-well or 394-well plate) and cheap (<1$ per assay).

Next, we are going to use this biosensor to test the functional domains of RBD interacting with ACE2. Previous studies indicate that the receptor binding motif (RBM) in the RBD (Fig. 2A) is more important for its interaction with ACE2^18^. We have made RBM-Lg construct and compared its activity with that of RBD *in vitro*. Although RBD-Lg and RBM-Lg were expressed at similar levels in cells (Fig. 2B), compared to RBD-Lg/Sm-ACE2, the biosensor activity of RBM-Lg/Sm-ACE2 *in vitro* is only 20% of that of RBD-Lg/Sm-ACE2, suggesting that other sequences downstream of RBM (aa. 541-685) are also important for its interaction with ACE2. Previous structural study shows that the peptidase domain or ectodomain (aa. 18-615) of ACE2 (Fig. 2D) is important for its interaction with RBD^8^. However, the minimum region required for the interaction of ACE2 with RBD has not been mapped. To map the minimum region of ACE2 critical for its interaction with RBD, we further truncated the protease domain (Fig. 2D). Our biosensor analysis shows that although similar protein levels were detected for Sm-ACE2-18-615 and Sm-ACE2-18-515 (Fig. 2E), the biosensor activity of Sm-ACE2-18-515/RBD-Lg was significantly lower than that of Sm-ACE2-18-615/RBD-Lg (Fig. 2F). Further ACE2 deletion (Sm-ACE2-18-415) stabilized the ACE2 protein (Fig. 2E) but reduces biosensor activity (Fig. 2F), suggesting that aa. 18-615 of ACE2 has the full-length RBD-binding domain. Based on structure of ACE2-RBD complex^14^, we also mutated 4 residues on ACE2 important for binding of ACE2 to RBD. Although similar levels of proteins were expressed for both WT and mutant ACE2 (Fig. 2G), *in vitro* luciferase assay shows that mutation of ACE2 K31 or M82 into alanine (A) significantly reduces biosensor activity (Fig. 2H). Furthermore, mutation of ACE2 K353 into alanine almost abolish biosensor activity (Fig. 2H). These findings not only confirm the specificity of our SRAE2-BS but also identify K353A as a critical residue for RBD:ACE2 PPI.

**Figure 2.**
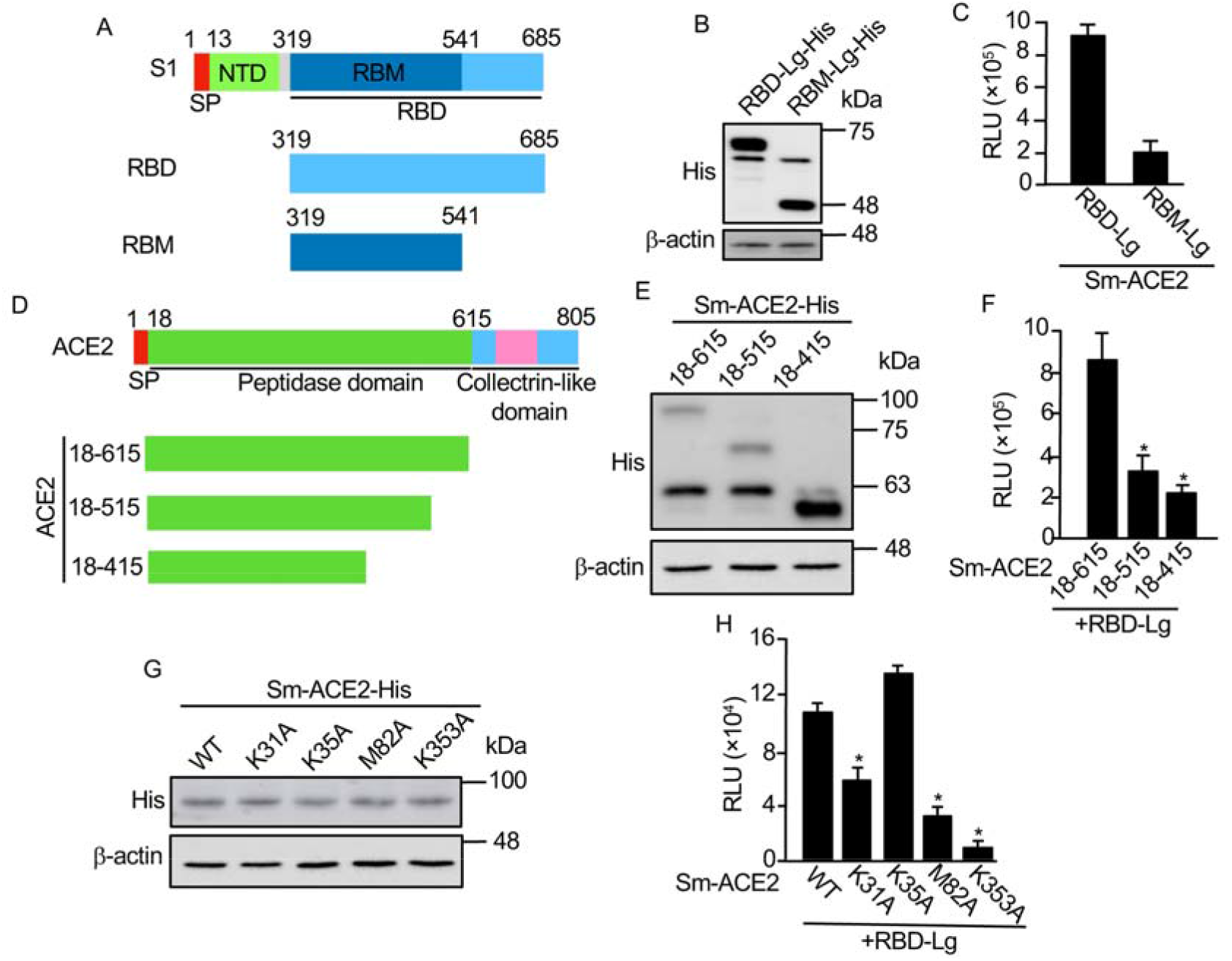
Mapping minimum region and critical residues for RBD:ACE2 PPI. **A.** Schematic of RBD deletions. RBD, receptor binding domain; SP, signal peptide; NTD, N-terminal domain; RBM, receptor binding motif. B, E, G. Western blot analysis of protein expression. C, F, H. *In vitro* luciferase assay. Equal amounts (1 μg) of each biosensor component were incubated together for 30 min, followed by luciferase assay; D. Schematic of ACE2 deletions. Experimental procedures and labels are as described in Figure 1. *, p<0.05

Currently, no SMs disrupting RBD:ACE2 PPI have been used for COVID-19 clinical treatments. Most studies predict candidate SMs inhibiting RBD:ACE2 PPI by using computer modeling^5,19,20^. We have used our SRAE2-BS to test several candidate SMs (i.e. Baicalin, Hesperidin, Sculellarin, Theaflavin, and Heparin) predicted by computer modeling to be able to disrupt RBD-ACE2 PPI^19–21^. Surprisingly, out of 5 SMs tested, only Theaflavin can significantly suppress SRAE2-BS activity with an IC50 of 41.7 μM (Fig. 3A). This result suggests that a functional analysis such as our biosensor assay is required to finally validate the true SMs disrupting RBD-ACE2 PPI. Monoclonal antibodies have previously been shown to be able to bind to SARS-CoV-2 RBD and disrupt RBD-ACE2 interaction, which inhibits viral entrance into cells^22^. We have used a VHHR72 anti-RBD antibody, which has previously been shown to effectively inhibit RBD-ACE2 PPI^10, 23^, to test our biosensor. The biosensor activity is suppressed by VHH72 rather than IgG control in a dose-dependent manner (Fig. 4B). We next synthesized peptides based on RBD-ACE2 complex structure and peptide-protein docking ranking^8, 14^ Although ACE2-P1 has no effect on SRAE2-BS activity, ACE2-P2 significantly suppresses SRAE2-BS with IC50 of 42 μM (Fig. 3C). Interestingly, different from ACE2-derived peptides targeting virus RBD, peptides RBD-P1, P2, P3 targeting ACE2 significantly suppressor SRAE2-BS activity in a dose-dependent manner (Fig. 3D) with IC50 of 9.0, 6.8, and 1.1 μM, respectively. Our findings clearly show that our SRAE2-BS can be a very useful and simple tool to quickly validate SMs, antibodies, and peptides disrupting RBD-ACE2 PPI for anti-COVID-19 therapy.

**Figure 3.**
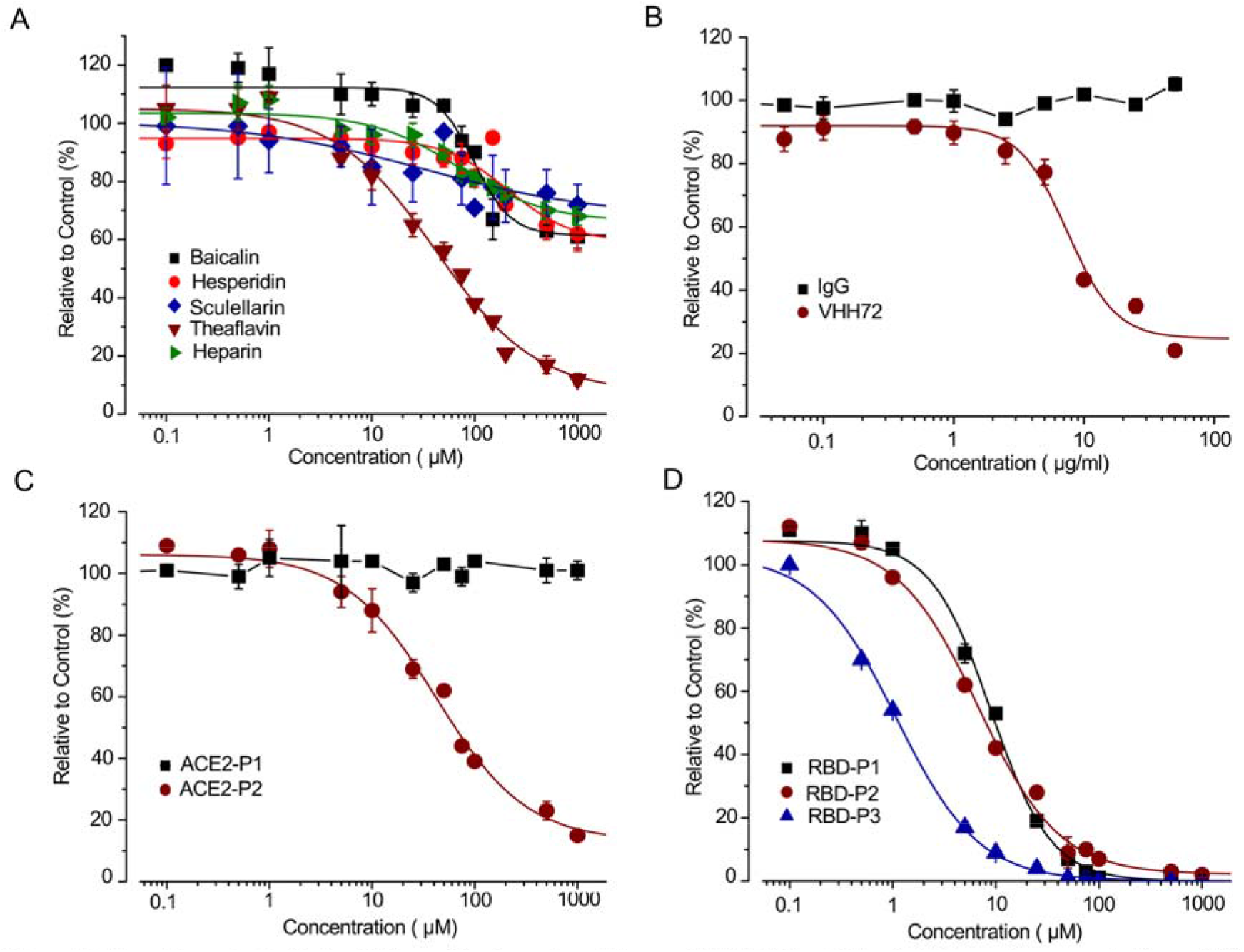
Dose-dependent effect of SMs, antibody and peptides on SRAE2-BS activity. A-C. Increasing concentrations of SMs (A), IgG (control) or VHH72 anti-RBD antibody (B), or ACE2-P1-P2 peptides (C) were preincubated with RBD-Lg-His, and subsequently incubated with Sm-ACE2-His for 30 min, followed by measurement of RLU. D. Increasing concentration of RBD-P1-P3 peptides were preincubated with Sm-ACE2-His, and subsequently incubated with RBD-Lg-His, followed by measure of RLU. The mean+/-standard error (S.E.) of relative values (relative to control) of triplicate samples were calculated in comparison to samples without treatment (control).

**Figure 4.**
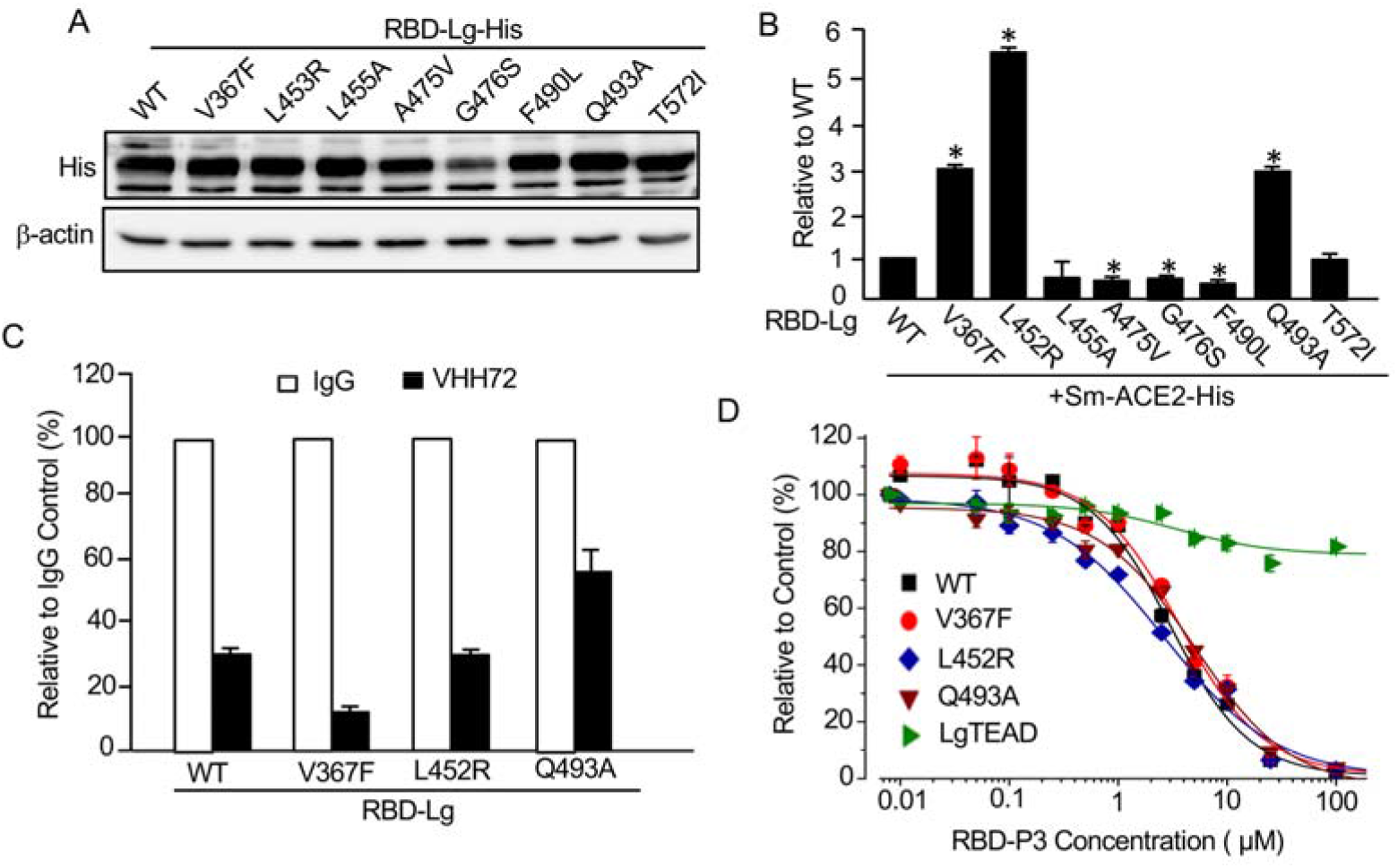
Mutations of SARS-CoV-2 RBD on its interaction with ACE2 and response to antibody and peptides. A. Western blot analysis of wild-type (WT) and mutant RBD-Lg-His. B. Mutations of RBD on its ACE2-binding ability. *In vitro* luciferase assay was carried out as described in Figure 1E. C. Mutations of RBD on their response to antibody. Protein lysates extracted from WT or mutant RBD-Lg-His were preincubated with 50 μg/ml of IgG or VHH72, and subsequently incubated with Sm-ACE2 lysate, followed by luciferase assay. D. Mutations of RBD on their response to peptide inhibition. Experimental procedures and data analysis are as described in Figure 3D.”, p<0.05

One character of coronavirus is the high frequency of mutations during spreading. Although the mutation rate for SARS-CoV-2 is not very high, many mutations have been detected in patients all over the world. However, since there is no system for rapid analysis of these mutations, how these mutations affect their binding to ACE2 is largely unknown. By taking the advantage of our newly constructed SRAE2-BS, we analyzed some of the RBD mutants detected in COVID-19 patients worldwide^24–29^. Except for RBD-G476S mutant, which is not stable in cells, compared to RBD-WT, western blot analysis indicates that the expression levels of RBD mutants are similar (Fig. 4A). When the same amounts of Sm-ACE2-His protein lysates were mixed with those of WT or each mutant RBD, RBD-L452A and RBD-T572I have similar affinity to Sm-ACE2 to RBD-WT, whereas A474V, G476S, and F490L mutants in RBD significantly reduce its ability to bind to ACE2 (Fig. 4B). Most interestingly, compared to WT RBD, V367F, L452R and Q493A RBD mutants obtained 3-5-fold increase in their bindings to ACE2 (Fig. 4B), suggesting that some of the RBD mutants indeed enhance the binding of virus to its receptor, therefore, probably resulting in higher entrance into cells and infectibility. Consistent to our findings, SARS-CoV-2 with V367F or L452R mutation was indeed shown to have enhanced entrance into cells, whereas virus with G476S, A475V or F490L mutations have reduced entrance into cells^28^. Our findings provide strong evidence that our SRAE2-BS is a surrogate assay for evaluating the ability of SARS-CoV-2 RBD mutants on ACE2 binding and virus infectibility. We next tested whether the mutants with enhanced binding to ACE2 are also sensitive to antibody suppression. As shown in Fig. 4C, although the binding of V367F and L453R mutants to ACE2 are still significantly suppressed by VHH72 at the same levels as WT RBD, whereas Q493A mutant is relatively insensitive to antibody suppression (Fig. 4C). We also tested whether the most potent peptide disrupting RBD-ACE2 PPI, RBD-P3, is able to suppress mutant RBD activity (Fig. 3D). Significantly, RBD-P3 can suppress the interaction of ACE2 with both WT and mutants at similar IC50 (Fig. 4D). As a control, the RBD-P3 peptide has little effect on LATS kinase biosensor, Sm-YAP/LgTEAD (Fig. 4D), suggesting that the effect of RBD-P3 on RBD:ACE2 PPI is specific. Together, these findings clearly suggest that our SRAE2-BS is a very useful tool for biochemical analysis of virus RBD mutants and their sensitivity to drug treatments.

Finally, we also used our biosensor to test whether it can evaluate the ability of antibodies in sera of convalescent COVID-19 patients in blocking RBD:ACE2 PPI. Since unknown molecules existing in sera of both healthy persons and COVID-19 patients interfere our biosensor analysis, we were unable to evaluate our biosensor using serum samples (data not shown). Further optimization for assay procedures and conditions, i.e., using affinity chromatography to extract antibodies of interest from the sample matrix, is required for future application of this biosensor for detecting the ability of antibodies produced in COVID19 patients in suppressing RBD-ACE2 PPIs.

## CONCLUSION

In this study, we have developed an ultrasensitive, simple, rapid biochemical biosensor tool to analyze the interaction of virus RBD and human ACE2 both in living cells and *in vitro*. It will have significant implications and applications in both basic research on SARS-CoV-2/ACE2 biology and COVID-19 clinical diagnosis and therapeutic drug discovery.

## Supporting Information

Table S1-S2.

## AUTHOR INFORMATION

### Author Contributions

X.Y., L.L., Y.T. W., and P.S. conceived of the biosensor and its applications. X.Y., LL., Y.H., Y.W.S., and S.S. carried out the experiments. D.Z. made figures. Y.G. collected and provided COVID-19 patient serum samples. X.Y. and L.L. wrote the manuscript. L.L., Y.T.W., Y.G., P.S., and D.Z. revised the manuscript.

## ACKNOWLEDGMENT

We thank Dr. Guojun Liu for inspiring discussion on the possible ways for COVID-19 diagnosis and therapies at the beginning of the COVID-19 pandemic, which made us start thinking about working on this project using our expertise.

## ABBREVIATIONS

aa: amino acid
ACE2: angiotensin-converting enzyme 2
DMEM: Dulbecco’s modified Eagle’s medium
HTP: high throughput
Lg/LgBiT: large BiT
RBD: receptor binding domain
PAGE: polyacrylamide gel electrophoresis
S: spike
SARS: severe acute respiratory syndrome coronavirus-2
SM: small molecule
Sm/SmBiT: small BiT
SPR: surface plasmon resonance
SLCA: split luciferase complementary assay
WT: wild-type

